# The ER tether VAPA is required for proper cell motility and anchors ER-PM contact sites to focal adhesions

**DOI:** 10.1101/2022.10.17.512434

**Authors:** Hugo Siegfried, Rémi Le Borgne, Catherine Durieu, Thaïs De Azevedo Laplace, Agathe Verraes, Lucien Daunas, Jean-Marc Verbavatz, Mélina L. Heuzé

## Abstract

Cell motility processes highly depend on the membrane distribution of Phosphoinositides (PInst), giving rise to cytoskeleton reshaping and membrane trafficking events. Membrane contact sites serve as platforms for lipid exchange and calcium fluxes between two organelles. Here, we show that VAPA, an ER membrane-resident contact site tether, plays a crucial role during cell motility. CaCo2 adenocarcinoma epithelial cells depleted for VAPA exhibit several collective and individual motility defects, disorganized actin cytoskeleton and altered protrusive activity. During migration, VAPA is required for the maintenance of PI(4,5)P2 levels at the plasma membrane, but not for PI(4)P homeostasis in the Golgi and endosomal compartments. Importantly, we show that VAPA regulates the dynamics of focal adhesions (FA) through its MSP domain, and is essential to stabilize and anchor ventral ER-PM contact sites to FA, thus mediating microtubule-dependent FA disassembly. To conclude, our results reveal unprecedented functions for VAPA-mediated membrane contact sites during cell motility and provides a dynamic picture of ER-PM contact sites connection with FA mediated by VAPA.

## INTRODUCTION

The lipid identity of membrane compartments is highly regulated within cells and plays a key role in a diversity of cellular processes. During cell motility, lipids, in particular Phosphoinositides (PInst), are local determinants of molecular events leading to cell polarization, the formation of actin-driven protrusions and turnover of focal adhesions (FA) (Hammond and Burke, 2020; Tsujita and Itoh, 2015). PI(4,5)P2, the most abundant PInst at the plasma membrane, controls FA dynamics by regulating the binding of talin to integrins (Martel et al., 2001; Thapa et al., 2012) and the polarized trafficking of integrins (Nader et al., 2016). PI(4,5)P2 is described as a modulator of actin cytoskeleton organization and dynamics, either directly through the recruitment of several actin-binding proteins (Senju et al., 2017), or indirectly by regulating the activation of RhoA (Lacalle et al., 2007; Xu et al., 2010) and Cdc42 GTPases (Daste et al., 2017). The product of PI(4,5)P2 phosphorylation, PI(3,4,5)P3, also plays a crucial role in cell motility at the plasma membrane (PM). Several i*n vitro* and *in vivo* studies have shown that PI(3,4,5)P3 acts as a “compass lipid” that stabilizes the direction of migration (Funamoto et al., 2002; Hannigan et al., 2002; Lam et al., 2012) through the activation of Rac1 (Kunisaki et al., 2006; Welch et al., 2002; Yoshii et al., 1999).

Lipids are synthesized in the endoplasmic reticulum (ER) and delivered to membranes through vesicular transport, but also through non-vesicular transport at so-named membrane contact sites (MCS) (Jackson et al., 2016). MCS are sites of close apposition between two membranes, often the ER membrane and the membrane of another organelle, at a distance of less than 80nm, where exchanges of lipids, Ca2+ and metabolites take place (Prinz et al., 2020; Scorrano et al., 2019; Wu et al., 2018). While the activity of lipid transfer at MCS was observed decades ago (Vance, 1990), their implication in patho-physiological processes such as cell motility has gained interest only recently (Prinz et al., 2020).

In this work, we studied the role of VAPA, a tethering protein at ER-mediated MCS during cell motility. VAPA is a member of the highly conserved VAP family of proteins, together with VAPB. VAP proteins are integral ER membrane proteins bridging the ER membrane to the membrane of other compartments by assembling with lipid transfer proteins that bind the target membrane, such as Nir proteins and OSBP-related (ORP) proteins. The assembly of VAP complexes relies on the interaction between the conserved N-terminal MSP (Major Sperm Protein) domain of VAPs and the FFAT (2 phenyl alanines in an acidic tract) motif on their partner. VAPs also contain a central coiled-coil dimerization domain and an ER transmembrane domain in their C-terminus (Murphy and Levine, 2016). VAP proteins are involved in several cellular functions such as lipid transport, membrane trafficking, the unfolded protein response pathway and microtubules organization (Kamemura and Chihara, 2019; Lev et al., 2008). VAP complexes with lipid transfer proteins control the homeostasis of PInst and sterols at various intracellular locations. At ER-PM contact sites, the VAP/Nir2 and VAP/ORP3 complexes regulate the transport of Phosphoinositol (PI) and PI(4)P respectively. Together with membrane-associated kinases and phosphatases, they contribute to modulating locally the pool of PI(4)P, PI(4,5)P2 and PI(3,4,5)P3 at the PM (Chang and Liou, 2015; Gulyás et al., 2020; Kim et al., 2013). In addition, the association of VAPs with ORP proteins modifies the level of PI(4)P at endosomal membranes and at the Golgi, therefore regulating trafficking events in these two compartments (Dong et al., 2016; Kawasaki et al., 2022; Mesmin et al., 2013). Moreover, VAP proteins mediate cholesterol transport at ER-Golgi, ER-endosome and ER-peroxisome MCS (Dong et al., 2019; Hua et al., 2017; Mesmin et al., 2013; Wilhelm et al., 2017). Depending on the cell type and organism, the loss of VAP proteins can lead to either a reduction (Peretti et al., 2008) or an accumulation (Mao et al., 2019) of PI(4)P levels in the Golgi membranes and its redistribution on endosomes (Dong et al., 2016). In yeast, depleting VAP orthologs results in the accumulation of PI(4)P at the plasma membrane (Stefan et al., 2011).

Altogether, these observations prompted us to hypothesize that VAPs, by regulating PInst homeostasis, could contribute to cell motility processes, which heavily rely on regulation of PM PInst. Importantly, two partners of VAPs at ER-PM contact sites, ORP3 and Nir2, have been shown to regulate cell motility processes through different pathways (D’Souza et al., 2020; Keinan et al., 2014; Lehto et al., 2008; Weber-Boyvat et al., 2015). As a first-line approach, we decided to investigate the effects of VAPA depletion alone on the motility of human adenocarcinoma Caco2 cells. We observed strong motility defects, suggesting that VAPA is required for proper cell motility and that VAPB is not sufficient to compensate for the loss of VAPA. We then aimed to understand the precise function of VAPA in this process. Here, we show that the depletion of VAPA strongly impacts the organization of the actin cytoskeleton, the dynamics of protrusions and the turnover of FA during migration. The role of VAPA stands mainly at the PM where it regulates PI(4,5)P2, the stability of ER-PM contact sites and their local anchoring at FA, thereby regulating FA disassembly.

## RESULTS

### VAPA is required for proper cell migration and cell spreading

To assess the role of VAPA during cell motility, we generated, using CRISPR/Cas9 gene editing, stable Caco-2 cells either depleted for VAPA (VAPA KO), or expressing non-targeting sequences (Control). VAPA KO cells exhibited a complete and specific depletion of VAPA protein (Fig.1A and 1B) without significantly affecting the levels of VAPB (which on the contrary had a tendency to increase), or its distribution in the ER (Sup Fig.1). We first assessed the capacity of VAPA KO and Control monolayers to migrate collectively upon space release on fibronectin-coated glass. In these conditions, VAPA KO monolayers filled the open space less efficiently than Control monolayers (Fig.1C and 1D, Sup Movie 1). The analysis of leader cells trajectories revealed that VAPA KO leader cells were displacing faster but with a remarkable lack of directionality compared to Control leader cells (Fig.1E and 1F), which could account for the slower colonization capacity VAPA KO monolayers. This collective migration impairment was at least partially cell-intrinsic, as single VAPA KO cells displacing on a 2D surface exhibited identical defects compared to VAPA KO leader cells in monolayers (Fig.1G and 1H). Moreover, upon plating, VAPA KO cells were spreading much faster than Control cells, also indicating a cell-autonomous defect arising in these cells (Fig.1I and 1J). Together, these results identify VAPA as a novel actor of collective and individual cell motility processes.

**Figure 1.**
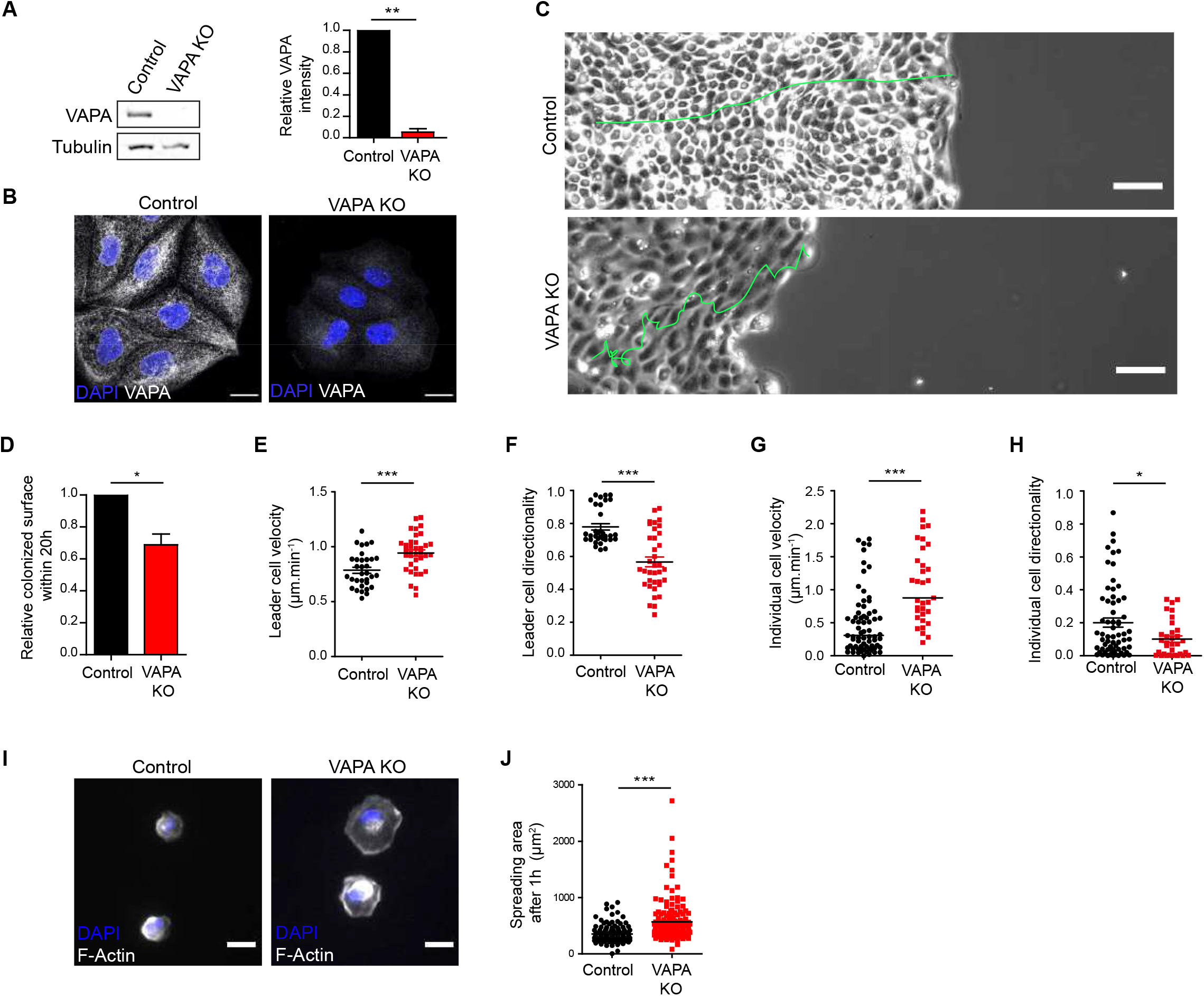
VAPA is required for proper cell migration and cell spreading. **A**. Left panel: Representative immunoblots showing the levels of VAPA and Tubulin in Control and VAPA KO cells. Tubulin expression level was used as loading control. Right panel: quantification of relative VAPA density in Control and VAPA KO cells normalized to Tubulin levels (mean±SEM from three independent experiments). **B**. Confocal images of Control and VAPA KO cells immunostained for VAPA. Scale bar 20 µm. **C**. Phase contrast images of Control and VAPA KO cells migrating collectively 48h after space release. The trajectories of a Control and VAPA KO leader cell over 20h are shown in green. Scale bar: 100 µm. **D**. Analysis of relative colonized surface within the last 20h by Control and VAPA KO cells (mean±SEM from three independent experiments). Data were analysed using a paired t-test. **E.F**. Analysis of Control and VAPA KO leader cell velocity (**E**) and directionality coefficient (**F**) calculated as the ratio between the net displacement and the trajectory length within the last 20h of collective migration (mean±SEM, Control: n=33 cells; VAPA KO: n=36 cells, from 3 independent experiments). **G.H**. Analysis of cell velocity (**G**) and directionality coefficient (**H**) of Control and VAPA KO individual cells displacing on fibronectin-coated glass during at least 3h (mean±SEM; Control: n=64 cells; VAPA KO: n=33 cells from 2 independent experiments). **I**. Epifluorescence images of Control and VAPA KO cells after 1 hour spreading on fibronectin-coated glass and stained as indicated. Scale bar: 20 µm. **J**. Analysis of Control and VAPA KO cells area after 1 hour spreading on fibronectin-coated glass (mean±SEM; Control: n=170 cells; VAPA KO: n=163 cells from 3 independent experiments). All data were analysed using non parametric Mann-Whitney t-test. (ns: non significant, ***P-values <0.001).

### VAPA regulates focal adhesions and actin cytoskeleton through its MSP domain

In order to characterize the motility defects of VAPA KO cells, we studied the organization of cell-matrix adhesions and actin cytoskeleton in these cells. To facilitate the analysis, we decided to focus on leader cells, standing at the leading edge of migrating monolayers, as these cells move in a given direction with a polarized and reproducible morphology. We observed an alteration of cell-matrix adhesions in VAPA KO cells that formed larger FA compared to Control cells (Fig.2A and 2D). Importantly, the expression of an exogenous wild-type form of VAPA in VAPA KO cells restored the size of FA, but it was not the case with the MSP-mutated form of VAPA (VAPA KDMD), indicating that VAPA regulates FA size through its association with proteins containing a FFAT motif (Fig.2A, 2D and Sup.Fig.2). We then compared the organization of actin cytoskeleton in both Control and VAPA KO cell lines. In Control cells, the F-actin and cortactin stainings revealed the presence of numerous and thick parallel transverse actin arcs at the front, accompanied by cortactin-rich subdomains of branched actin standing at the front edge corresponding to lamellipodial extensions (Fig.2B and 2C). We indeed observed periodic waves of protrusion-retraction arising in these cells every 15 minutes in average (Fig.2C and 2G). By contrast, VAPA KO cells exhibited disorganized and thinner transverse actin arcs, wider cortactin-rich subdomains occupying most of the leading edge and giving rise to long-lasting protrusions (Fig.2B,2C,2E-2G). Similarly to the FA phenotype, we were able to recover the size of cortactin-rich domains by expressing a wild-type form of VAPA but not the MSP mutant VAPA KDMD (Fig.2E). To conclude, we show that VAPA controls different aspects of cell motility, namely the maintenance of FA size and the preservation of the spatial and temporal organization of the actin cytoskeleton. These functions depend on the association of VAPA with FFAT-motif containing proteins such as lipid transfer proteins, suggesting that lipid transfer is probably required.

**Figure 2.**
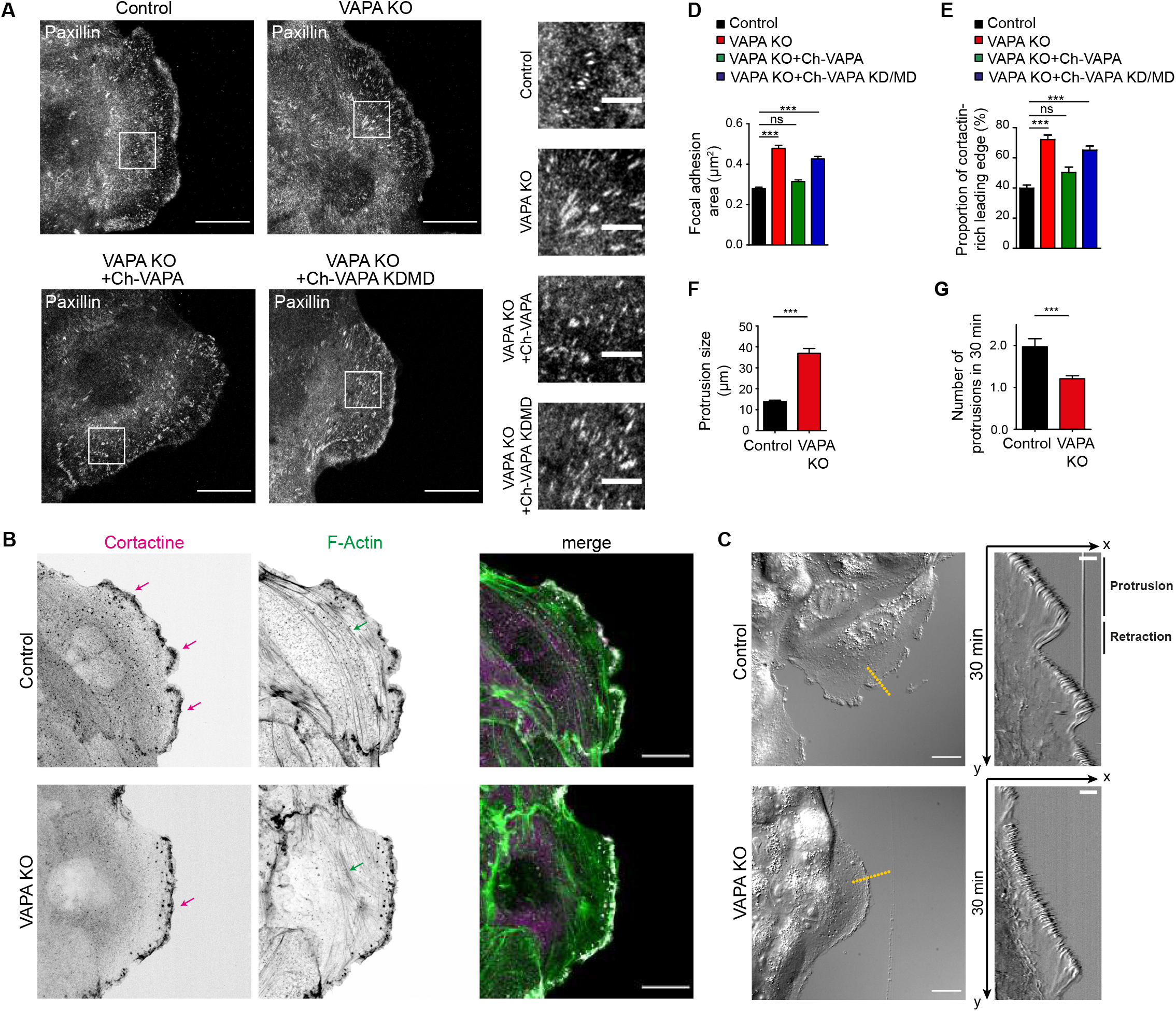
VAPA regulates focal adhesions and actin cytoskeleton through its MSP domain. **A**. Confocal images with zoom boxes of paxillin-labeled focal adhesions in leader cells from Control, VAPA KO, VAPA KO stably expressing wild-type mCherry-VAPA, or mCherry-VAPA KD/MD cell lines. Scale bar: 20 µm (5 µm in insets). **B**. Confocal images of actin cytoskeleton network in Control and VAPA KO leader cells stained for cortactin and F-actin. Magenta arrows point to cortactin-enriched domains at the leading edge. Green arrows point to F-Actin transversal arcs. Scale bar: 20 µm. **C**. Differential Interference Contrast (DIC) images (left) of Control and VAPA KO leader cells and kymographs (right) along the yellow lines from a 30 min movie at 1 frame every 3 seconds, showing protrusion and retraction phases of the leading edge. Scale bar: 20 µm (left) and 5 µm (right). **D**. Analysis of focal adhesion area quantified from images in A, in Control (n=2968 focal adhesions from 20 cells), VAPA KO (n=4329 focal adhesions from 22 cells), VAPA KO expressing mCherry-VAPA, (n= 3818 focal adhesions from 24 cells) and VAPA KO expressing mCherry-VAPA KD/MD (n=2801 focal adhesions from 19 cells) leader cells, from 3 independent experiments **E**. Analysis of leading edge proportion enriched with cortactin staining quantified from images in B, in Control, VAPA KO, VAPA KO expressing wild-type mCherry-VAPA and VAPA KO expressing mCherry-VAPA KD/MD leader cells, (mean±SEM; n=29 cells for each cell line, from 3 independent experiments). **F**. Analysis of the size of cortactin-enriched domains at the leading edge, quantified from images in B, in Control and VAPA KO leader cells (mean±SEM; Control n= 77 cells; VAPA KO n=72 cells, from 3 independent experiments) **G**. Quantification of protrusion phases frequency per 30min quantified from the kymographs in C, in Control and VAPA KO leader cells (mean±SEM; Control n=16 cells; VAPA KO: n=21 cells from 3 independent experiments). All data were analysed using non parametric Mann-Whitney t-test (ns: non significant, ***P-values <0.001).

### VAPA controls PI(4,5)P2 levels at the PM, but is not essential for PI(4)P homeostasis in the Golgi and endosomal compartments

We then intended to decipher the mechanisms by which VAPA could regulate FA size, the actin cytoskeleton and cell motility. To test the lipid transfer hypothesis, we first characterized to which extend its deletion affected PInst homeostasis. Previous studies have shown that depleting both VAPA and VAPB affected the levels of PI(4)P at the Golgi and PM (Mao et al., 2019; Stefan et al., 2011), and induced the accumulation of PI(4)P in early endosomes (Dong et al., 2016). Using a GFP-PH-OSBP probe, we detected a major intra-cellular pool of PI(4)P at similar levels in the Golgi compartment in Control and VAPA KO cells (Fig.3A-C). Importantly, the depletion of VAPA alone was not sufficient to observe an accumulation of PI(4)P in early endosomes (Fig.3A-C). This result suggests that VAPB can compensate for the absence of VAPA at ER-Golgi and ER-endosome contact sites thus maintaining PI(4)P homeostasis at the Golgi and endosomes in VAPA KO Caco-2 cells. However, at the PM, the depletion of VAPA induced a decrease of PI(4,5)P2 which is the most abundant PInst in this membrane compartment (Fig.3D-3G). This decrease was detected at the dorsal side of cells by immunofluorescence staining (Fig.3D and 3F), and at the protrusive cortactin-rich leading edge of migrating cells by measuring the PM/cytoplasm ratio of RFP-PH-PLCδ1 probe for PI(4,5)P2 (Fig.3E and G). Altogether, these results show that in Caco-2 cells, VAPA is required to maintain a certain level of PI(4,5)P2 specifically at the plasma membrane, but is dispensable for PI(4)P homeostasis in the Golgi and endosomal compartments.

**Figure 3.**
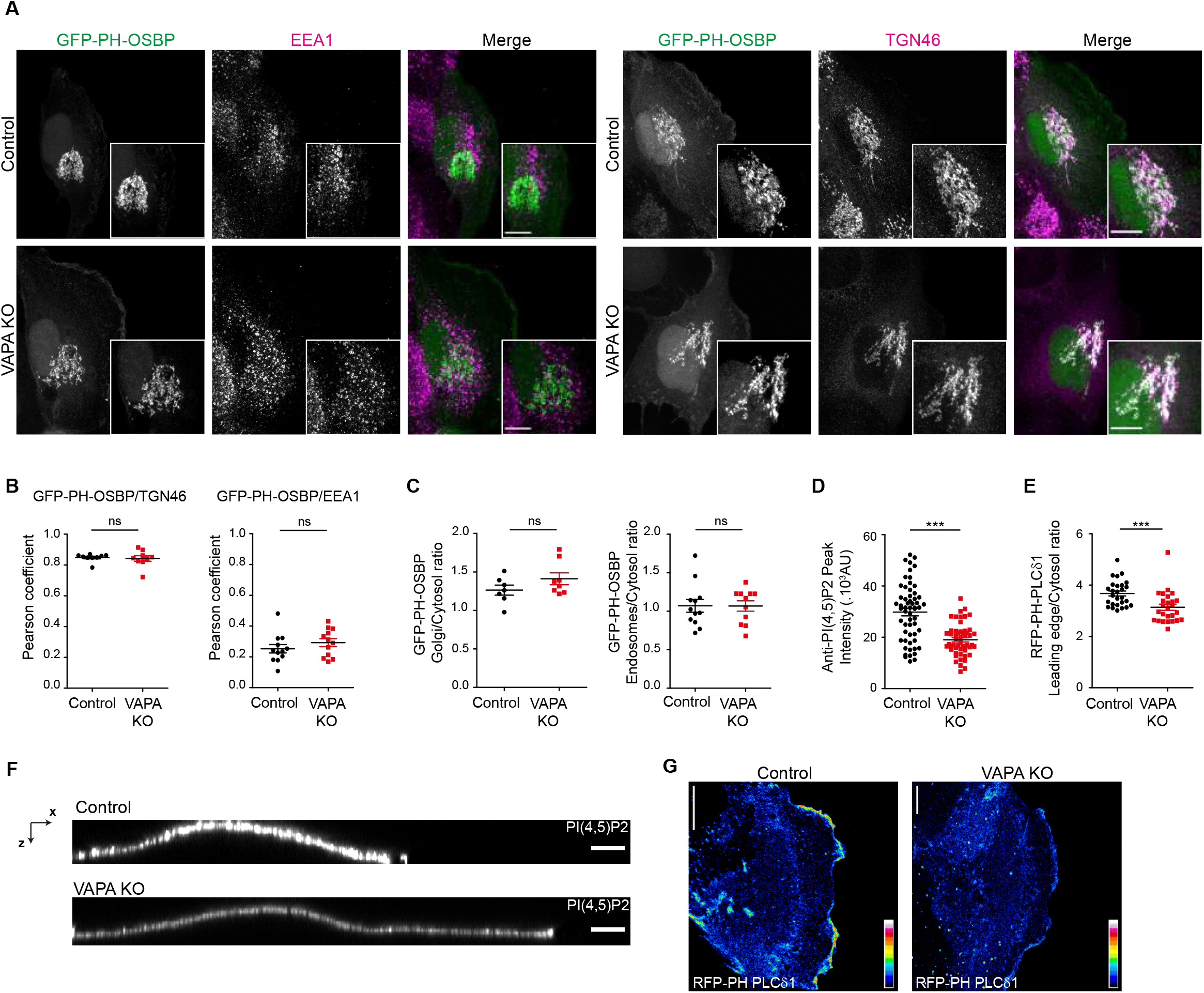
VAPA controls PInst homeostasis at the PM, but not in other compartments. Confocal images and zoom boxes of PI(4)P distribution in Control and VAPA KO leader cells expressing GFP-PH-OSBP and immunostained for EEA1 or TGN46. Scale Bar: 10 µm. Analysis of colocalization by Pearson coefficient determination between GFP-PH-OSBP and EEA1 or TGN46 signals, quantified from images in A, in Control and VAPA KO leader cells (mean±SEM; Control: n=9 cells and 12 cells respectively; VAPA KO: n=9 cells and 12 cells respectively, from 3 independent experiments). **C**. Analysis of Golgi(TGN46)/cytosol (left) and Endosomes(EEA1)/Cytosol (right) ratio of GFP-PH-OSBP signal, quantified from images in A, in Control and VAPA KO leader cells (mean ±SEM; Control: n=12 cells; VAPA KO: n=11 cells, from 3 independent experiments). **D**. Analysis of RFP-PH-PLCδ1 peak intensity, quantified from images in F, in Control and VAPA KO leader cells (mean±SEM; Control: n=58 cells; VAPA KO: n=53 cells, from 3 independent experiments). **E**. Analysis of PM/Cytosol ratio of RFP-PH-PLCδ1, quantified from images in G, at the leading edge of Control and VAPA KO leader cells (mean±SEM; Control: n=27 cells; VAPA KO: n=26 cells, from 3 independent experiments). **F**. XZ view of confocal images of Control and VAPA KO cells in cluster immunostained for PI(4,5)P2. Scale Bar: 5 µm. **G**. Confocal images of PI(4,5)P2 distribution in Control and VAPA KO leader cells expressing RFP-PH-PLCδ1, represented as a color-coded heat map. Scale bar: 10 µm. All data were analysed using non parametric Mann-Whitney t-test (ns: non significant, ***P-values <0.001).

### VAPA stabilizes ventral ER-PM contact sites at the front of migrating cells

Based on our observation pointing to a specific role of VAPA for the PI(4,5)P2 homeostasis at the PM, we questioned the spatio-temporal distribution of ER-PM contacts sites during cell migration and the role of VAPA in this organization. To this aim, we took advantage of a fluorescent probe, GFP-MAPPER, which selectively labels ER-PM contact sites (Chang et al., 2013). When expressed in Caco-2 cells, GFP-MAPPER distributed as puncta along ER tubules (Fig.4A) and appeared at ER-PM appositions detected by TIRF microscopy at the cell bottom (Fig.4B and Sup Movie 2). At the front of leader cells, GFP-MAPPER puncta divided in two subpools: the dorsal subpool ongoing forward movement with the migrating cell and the ventral subpool remaining immobile relative to the substrate, suggesting that ventral ER-PM contact sites might be anchored to cell-matrix adhesions (Fig.4C). Using electron microscopy, we were indeed able to identify numerous ER-PM contact sites sitting at the ventral leading edge of migrating cells (Fig.4D). At the ultrastructural level, the depletion of VAPA had no significant effect on ER-PM contact sites regardless of their location (Sup Fig.3). However, the dynamic analysis of GFP-MAPPER puncta revealed that ventral ER-PM contact sites, which barely moved (400nm/min on average) and persisted generally more than 10 minutes in Control cells, were significantly less stable both spatially and temporally in VAPA KO cells (Fig.4E-G and Sup Movie 3). To conclude, our results identify a sub-pool of ER-PM contact sites docked to the substrate at the front of migrating cells, and whose spatio-temporal stability requires VAPA. Together with the fact that FA size is altered in VAPA KO cells, these results prompted us to hypothesize that VAPA might accomplish a function at FA.

**Figure 4.**
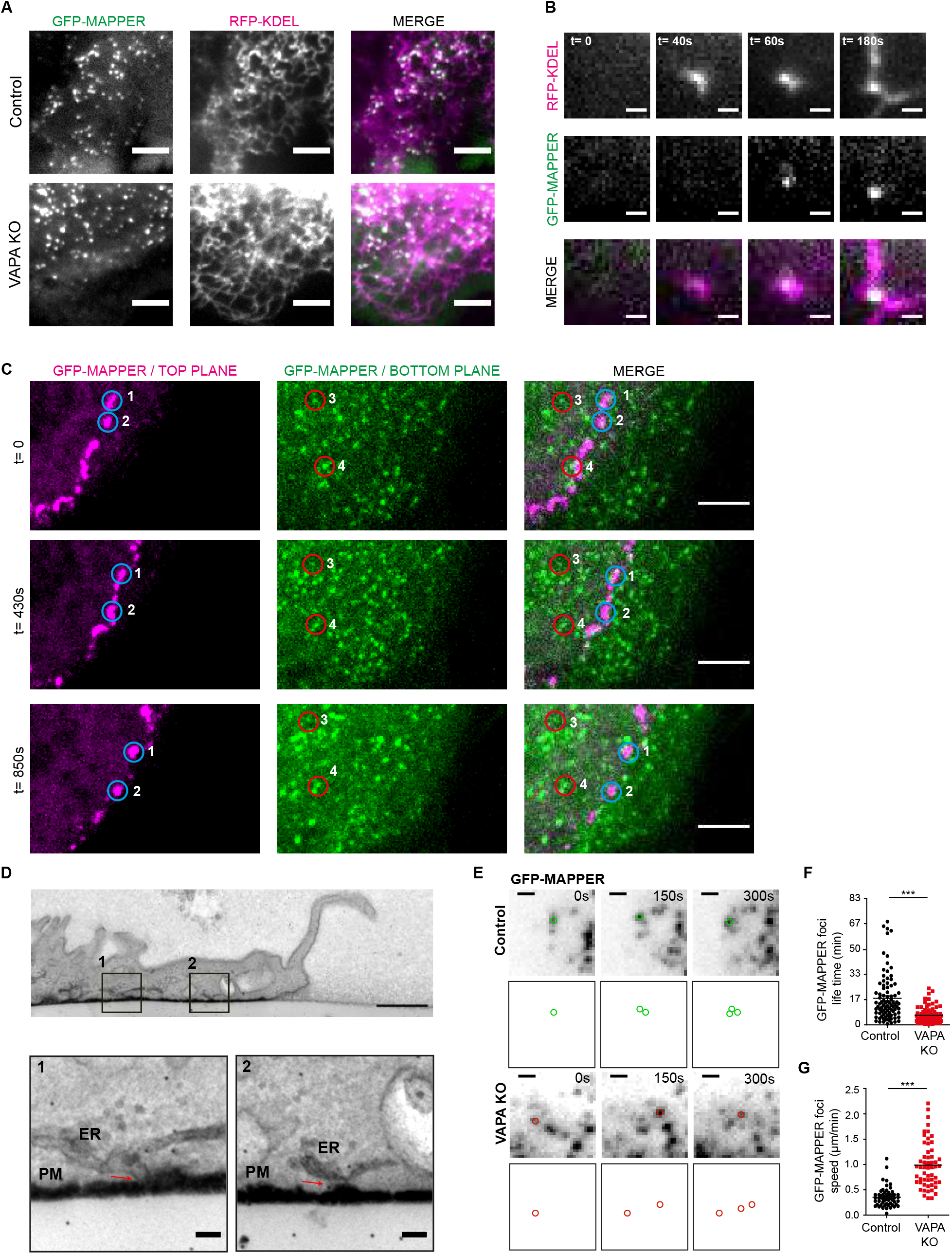
VAPA stabilizes ventral ER-PM contact sites at the front of migrating cells. **A**. TIRF microscopy images of GFP-MAPPER foci distribution along the ER in Control and VAPA KO leader cells expressing GFP-MAPPER and RFP-KDEL. Scale bar: 5 µm. **B**. Sequential TIRF microscopy images of ER and GFP-MAPPER foci accumulation at the front of a Control leader cell expressing GFP-MAPPER and RFP-KDEL. Scale bar: 1 µm. **C**. Sequential spinning disk images of dorsal (magenta) and ventral (green) GFP-MAPPER foci at the front of a Control leader cell expressing GFP-MAPPER. Top and bottom plane images were color-coded in magenta and green respectively. Circles highlight single GFP-MAPPER foci at the dorsal (blue) and ventral (red) sides. Scale bar: 5 µm. **D**. Transmission Electron Microscopy images of transversal cuts of a Control leader cell, showing the leading edge. Arrows point to ER-PM contact sites at the bottom of the cell. Scale bar: 1 µm (top) and 100 nm in insets 1 and 2. **E**. Sequential TIRF microscopy images of GFP-MAPPER foci at the front of Control and VAPA KO leader cells expressing GFP-MAPPER. Individual GFP-MAPPER foci are pictured in the frames below images. Scale bar: 1 µm. **F.G**. Analysis of the lifetime (**F**) and the speed (**G**) of ventral GFP-MAPPER foci, quantified from images in E, in Control and VAPA KO leader cells (mean±SEM; Control: n=8 cells; VAPA KO: n=6 cells; at least 10 ER-PM MCS were analysed per cell, from 4 independent experiments). All data were analysed using non parametric Mann-Whitney t-test (***P-values <0.001).

### VAPA promotes microtubule-dependent FA disassembly

To further investigate the role of VAPA in FA turnover, we studied more precisely FA dynamics in Control and VAPA KO cells during cell migration by TIRF microscopy. While the assembly rate of FA was similar in both cell lines, their disassembly rate was slower in VAPA KO cells (Fig.5A, 5B and 5E). In addition, VAPA KO FA exhibited a longer lifetime in average (Fig.5C and 5E), with a majority lasting at least 45 minutes (Fig.5D and E). To test whether VAPA-mediated disassembly of FA relied on the microtubule network, we synchronized FA disassembly by nocodazole washout experiment in migrating cells and measured the evolution of FA size, as described previously (Ezratty et al., 2005). Our results clearly show that VAPA KO FA failed to disassemble upon microtubule recovery, contrary to Control FA that lost half of their size within 15 minutes after nocodazole washout (Fig.5F and 5G). These results demonstrate that VAPA is required for microtubule-dependent FA disassembly during migration of Caco2 cells.

**Figure 5.**
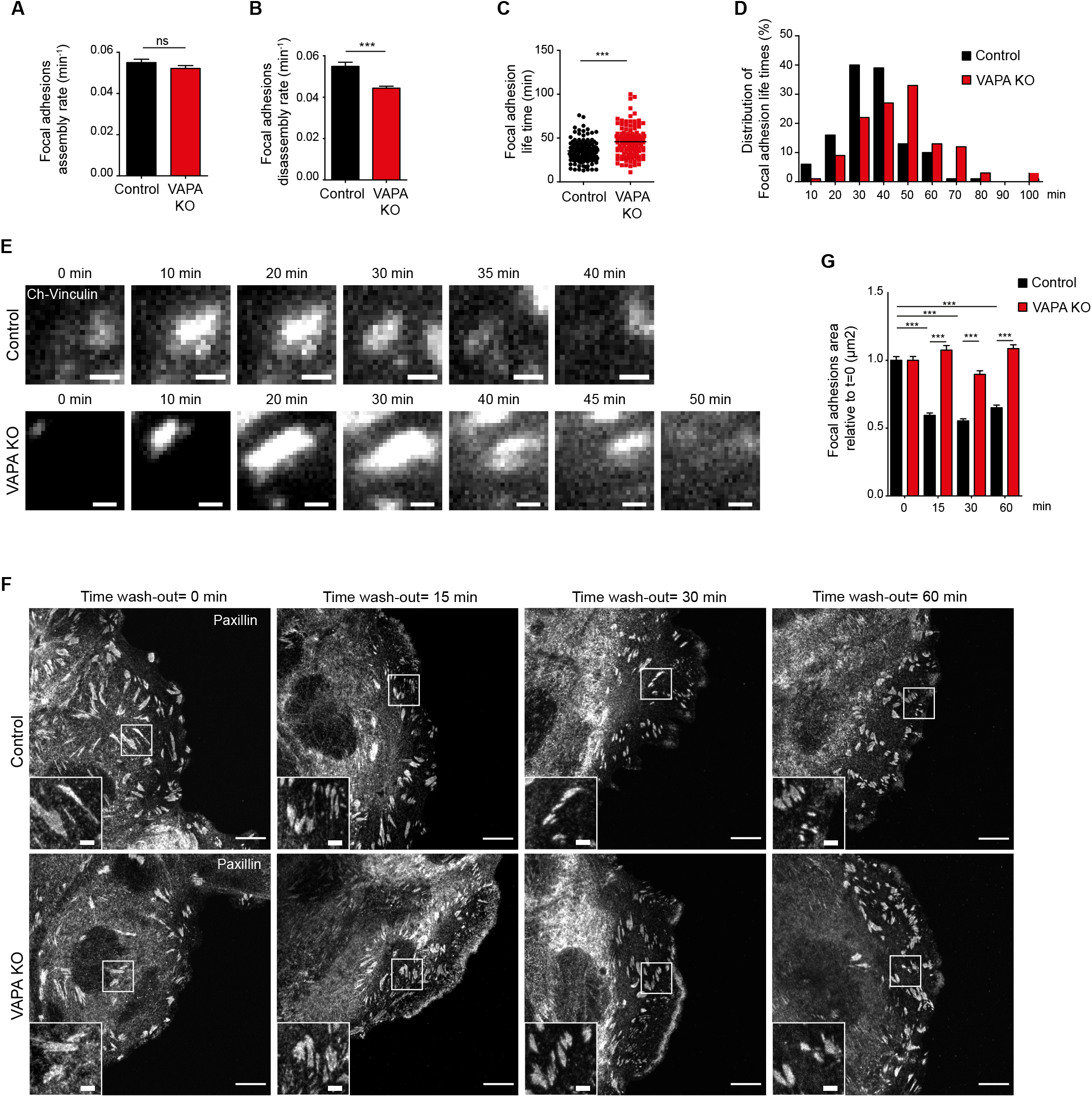
VAPA promotes microtubule-dependent FA disassembly. **A-C** Analysis of assembly rate (**A**), disassembly rate (**B**) and lifetime (**C**) of focal adhesions (FA), quantified from time-lapse images in E, in Control and VAPA KO leader cells (mean±SEM; Panels (A) and (B): Control: n=217 FA from 9 cells; VAPA KO: n=342 FA from 8 cells, from 3 independent experiments. Panel (C): Control: n=133 FA from 9 cells; VAPA KO: n=126 FA from 8 cells, from 3 independent experiments). **D**. Distribution of focal adhesion (FA) life times, quantified from time-lapse images in E, in Control and VAPA KO leader cells (Control: n=133 FA from 9 cells; VAPA KO: n=126 FA from 8 cells, from 3 independent experiments). **E**. Sequential TIRF microscopy images of focal adhesions in Control and VAPA KO leader cells expressing mCherry-Vinculin. Scale Bar: 1 µm. **F**. Sequential Confocal images of focal adhesions after nocodazole treatment and wash-out in migrating Control and VAPA KO leader cells immunostained for paxillin. Scale bar: 10 µm (2 µm in insets). **G** Analysis of relative focal adhesions (FA) size, quantified from images in F, in Control and VAPA KO leader cells after 0 min, 15 min, 30 min and 60 min after nocodazole wash-out (mean±SEM. FA from 22 to 25 cells were analysed, from 3 independent experiments. Control T0min: n=6053 FA; Control T15 min: n=5146 FA; Control T30 min: n=5543 FA; Control T60 min: n=3913 FA; VAPA KO T0 min: n=4481 FA, VAPA KO T15 min: n=3878 FA, VAPA KO T30 min: n= 4165 FA ; T60 min : n=4889 FA). All data were analysed using non parametric Mann-Whitney t-test (ns: non significant, ***P-values <0.001).

### VAPA mediates the anchoring of ER-PM contact sites to FA

To understand the function of VAPA at FA, we analysed the repartition and the dynamic of ER-PM contact sites relative to FA by TIRF microscopy. In Control cells, the majority of vinculin-stained FA (80%) established a contact with ventral ER-PM contact sites whereas it was the case for only 30-40% of FA in VAPA KO cells (Fig.6A and 6B). We then monitored the time course of mCherry-Vinculin and GFP-MAPPER signals within the perimeter of single FA (Fig.6C-D and Sup Movie 4). GFP-MAPPER foci started to accumulate in the FA vicinity after its assembly, suggesting that ER-PM contact sites were anchoring to preexisting FA. Importantly, the disassembly of FA started immediately upon GFP-MAPPER recruitment and evolved linearly in parallel to GFP-MAPPER accumulation (Fig.6E and Sup Movie 4). Thus, we show that VAPA is required for the anchoring of ER-PM contact sites to FA, which happens at the time of FA disassembly.

**Figure 6.**
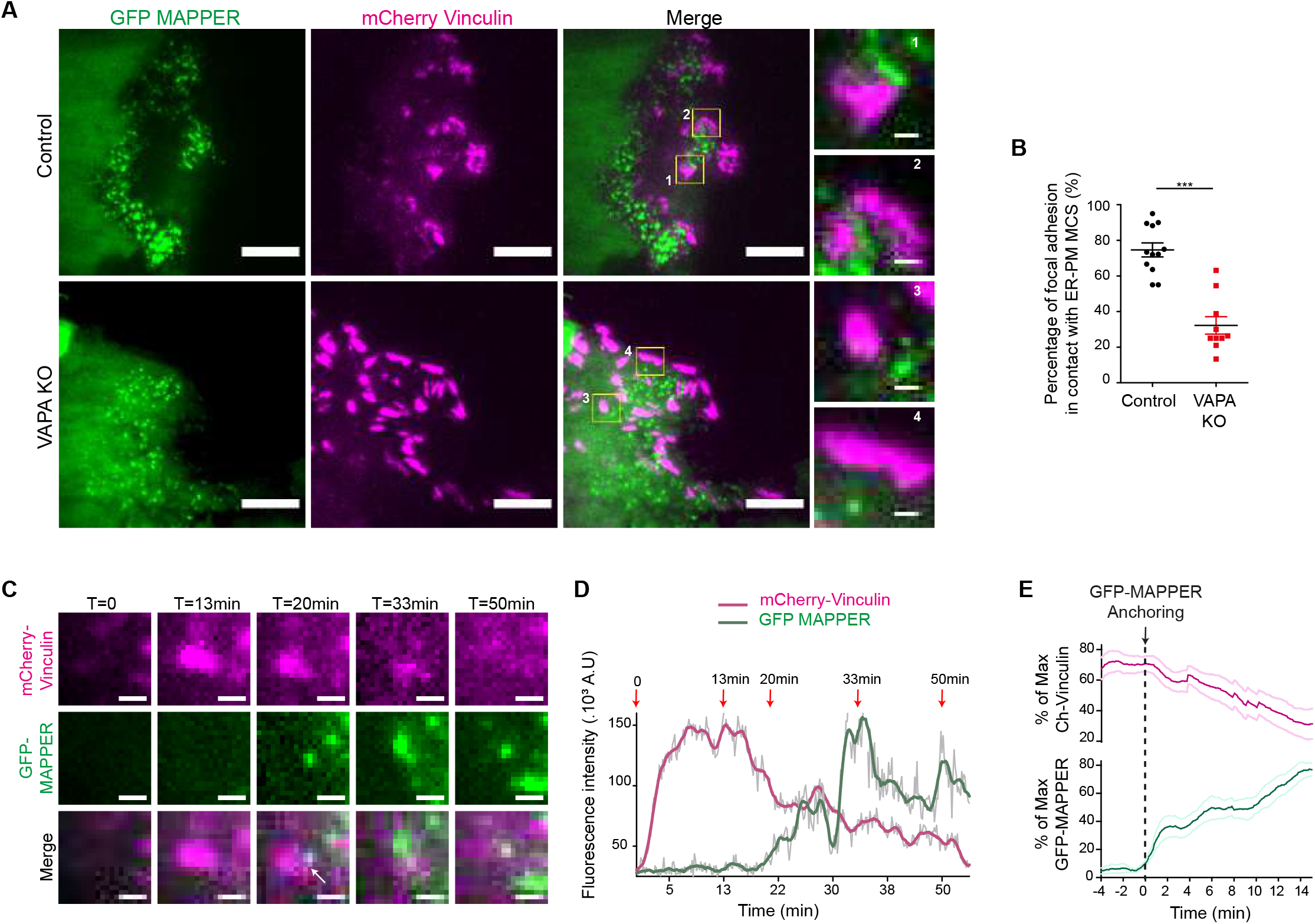
VAPA mediates the anchoring of ER-PM contact sites to FA. A. TIRF microscopy images of ER-PM contact sites and focal adhesions in Control and VAPA KO leader cells expressing GFP-MAPPER and mCherry-Vinculin. Scale bars: 10 µm (1 µm in insets). **B**. Analysis of the percentage of focal adhesions in contact with ER-PM contact sites, quantified from images in A, in Control and VAPA KO leader cells (mean±SEM; Control: n=12 cells; VAPA KO: n=10 cells, from 3 independent experiments). **C**. Sequential TIRF microscopy images of ER-PM contact sites and focal adhesions in a Control leader cell expressing GFP-MAPPER and mCherry-Vinculin. Scale bar: 1 µm. **D**. Time course of GFP-MAPPER (green) and mCherry-Vinculin (magenta) signals during the lifetime of the focal adhesion pointed in C, within a region that comprises the entire FA + 2 pixels. **E**. Time course of mCherry-Vinculin and GFP-MAPPER signals obtained by TIRF microscopy in Control leader cells, as in C., starting 4 minutes before the accumulation of GFP-MAPPER in FA regions (comprising the entire FA + 2 pixels). The signals were all smoothed, readjusted to the minimal value and expressed as % of the maximal value (mean±SEM; n=14 FA from 5 cells, from 3 independent experiments). All data were analysed using non parametric Mann-Whitney t-test (***P-values <0.001).

## DISCUSSION

In this work, we identify VAPA as a crucial player in cell motility, with implications in different aspects of this process. We show that VAPA is required for both single and collective cell migration, cell spreading, protrusive waves, actin cytoskeleton dynamics and FA turnover. Our study reveals a non-redundant function for VAPA in cell motility among the VAP family of proteins in Caco2 cells. Indeed, the presence of VAPB in VAPA KO cells was not sufficient to compensate the loss of VAPA. The function of VAPA in cell migration seems to arise from its specific role at ER-PM contact sites where it was necessary to maintain high levels of PI(4,5)P2 independently of VAPB. It was not the case at the Golgi or endosomes, where the depletion of VAPA alone had no effect on PI(4)P levels, which is in agreement with previous observations by Dong et al in VAPA/VAPB double knock-out cells re-expressing VAPB (Dong et al., 2016). This suggests that VAPB might compensate for the absence of VAPA at ER-Golgi and ER-endosomes contact sites and that migration defects in VAPA KO cells were not a result of a dysfunction in the secretory pathway. Together with previous studies (Dong et al., 2016; Mao et al., 2019; Peretti et al., 2008; Stefan et al., 2011), our results converge to the idea that VAPA and VAPB might fulfill both redundant and specific tasks in lipid homeostasis and cellular functions.

PI(4,5)P2 and its phosphorylation product PI(3,4,5)P3 have been described for decades as central orchestrators of cell motility through regulation of actin dynamics, FA maturation and turnover, membrane organization and curvature, and intracellular trafficking (Mandal, 2020; Tsujita and Itoh, 2015). Thus, the phenotypes that we observed in VAPA KO cells – namely disorganization of actin, protrusion defects, FA persistence, loss of directional movement and faster speed– could be solely explained by a global reduction in PI(4,5)P2 levels at the PM. However, our results clearly show that VAPA not only regulates PI(4,5)P2 homeostasis at the PM, but also fulfills a function at FA where it anchors ER-PM contact sites and favours their stability, thereby promoting microtubule-dependent FA disassembly. Importantly, we provide for the first time a quantitative spatio-temporal connection between ER-PM contact sites and FA dynamics in polarized and migrating cells. Based on these results, we propose a hypothetical model in which VAPA, through local regulation of lipid transfer in the vicinity of FA would induce the internalization of integrins through clathrin-mediated endocytosis which depends on PI(4,5)P2 levels (Chao et al., 2010; Nader et al., 2016) and on microtubule polymerization (Ezratty et al., 2009, 2005) (Fig.7). Interestingly, MCS have already been shown to directly regulate endocytosis in yeast (Encinar del Dedo et al., 2017).

**Figure 7.**
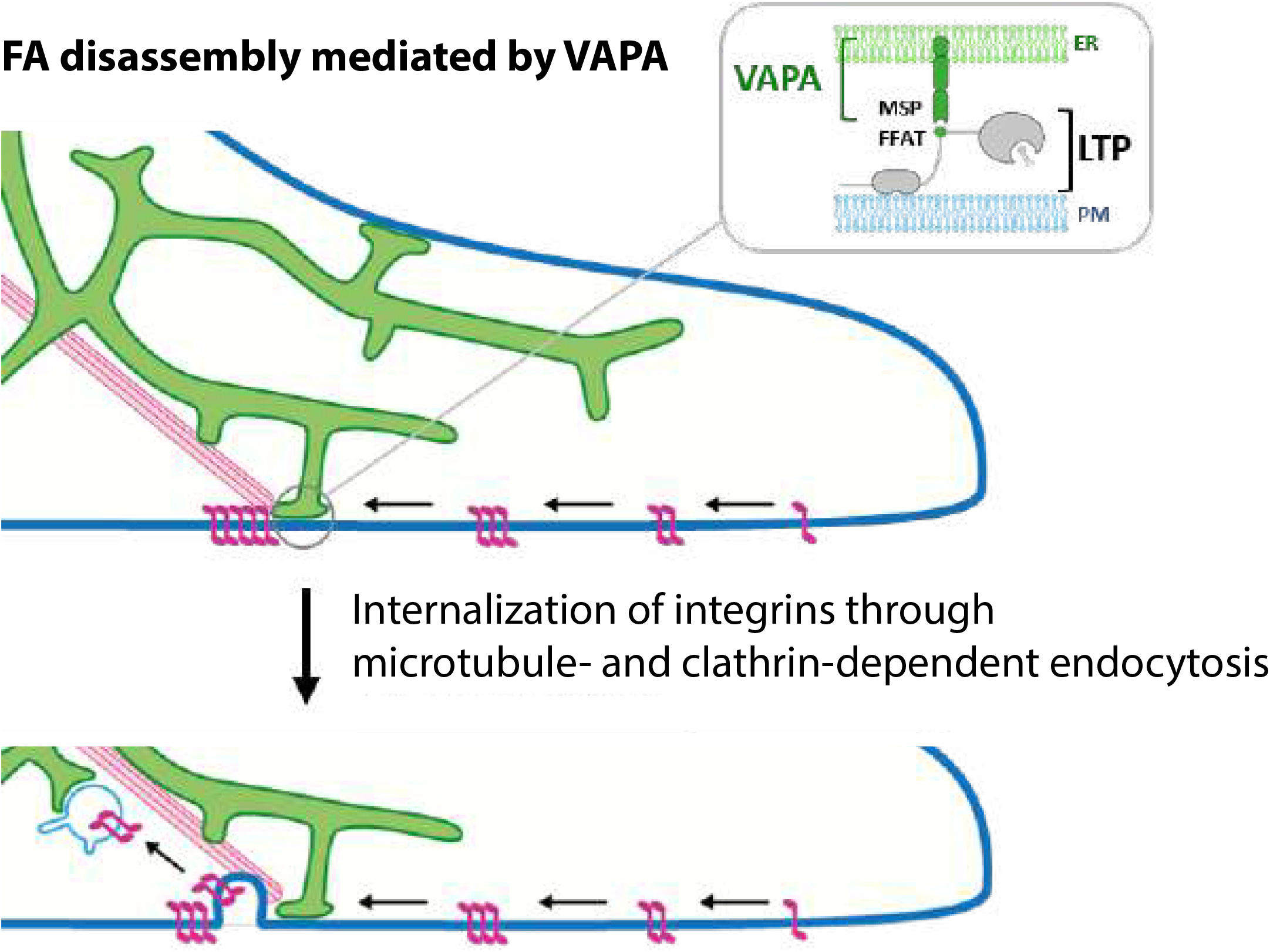
Hypothetic model depicting the role of VAPA in FA disassembly, based on our observations and previously published data (see Discussion part).

Which proteins could control the anchoring of VAPA-mediated ER-PM contact sites to FA? Previous studies have highlighted the role of ER proteins in FA turnover and maturation that could be potential candidates. Kinectin, an integral ER membrane protein interacting with kinesin, has been shown to control the apposition of ER to FA through a microtubule-dependent transport (Guadagno et al., 2020; Ng et al., 2016). The phosphatase PTP1B, which localizes to the cytoplasmic face of the ER and was identified as a partner of VAPA using a BioID approach (Antonicka et al., 2020), is targeted to newly formed FA and contributes to their maturation through dephosphorylation of its substrate p130Cas (Dadke and Chernoff, 2002; Hernández et al., 2006; Liu et al., 1998). In addition, we demonstrate that the function of VAPA at FA depends on its MSP domain of interaction with proteins containing a FFAT domain. Importantly, two lipid transfer proteins interacting with VAPA at ER-PM contact sites, Nir2 and ORP3, have been previously described as regulators of cell motility, establishing them as potential drivers for the specific function of VAPA at FA. Nir2, which interacts with VAPs at ER-PM and ER-Golgi contact sites and regulates PI transport (Chang and Liou, 2015; Kim et al., 2013; Peretti et al., 2008), favors epithelial-mesenchymal transition and tumor metastasis (Keinan et al., 2014). Similarly to VAPA KO cells, the depletion of Nir2 in HeLa cells induces a decrease of PI(4,5)P2 levels at the PM (Kim et al., 2013), indicating that a part of VAPA effect on PInst homeostasis could be supported by VAPA/Nir2-mediated lipid transfer. ORP3, which extracts PI(4)P from the PM at ER-PM contact sites (Gulyás et al., 2020), positively regulates the R-Ras pathway promoting cell-matrix adhesion (Lehto et al., 2008; Weber-Boyvat et al., 2015). More recently, ORP3 was found to be recruited to disassembling FA upon calcium influx and to favour their turnover through lipid exchange. Interestingly, the authors highlight a crosstalk between ORP3-mediated lipid exchange and STIM1/Orai1-mediated calcium influx at FA (D’Souza et al., 2020). Therefore, the role of VAPA-mediated contact sites at FA could not only be the local modulation of lipid transfer, but also the establishment of a common platform with STIM1, anchored to FA at ER-PM contact sites controlling lipid and calcium exchanges and driving FA maturation and turnover.

To conclude, our study reveals unprecedented functions of VAPA in cell motility processes and in local anchoring of ER-PM contact sites to FA. For a long time, we have assumed that PInst regulation at FA will principally be mediated by the interplay of lipid kinases and phosphatases. Our work reinforces the idea that lipid transfer would be an additional pathway for lipids to be controlled at FA and provides evidences for a spatio-temporal connection between MCS and FA mediated by VAPA. Finally, this work opens new roads to decipher the precise molecular determinants of VAPA function and to explore the role of VAPA during cancerogenesis.

## MATERIALS AND METHODS

### Cell culture

CaCo-2 cells, originally acquired from ATCC, were kindly provided by Dr. D. Delacour (Institut Jacques Monod, Paris). CaCo-2 CRISPR control and VAPA knock-out cell lines were maintained in culture in DMEM (1x) GlutMax 5g/L D-glucose + pyruvate medium (Gibco) supplemented with 20% fetal bovine serum (Gibco), 100 units/mL penicillin, 100 µg/mL streptomycin and 2 µg/Ml (Gibco) puromycin in 5% CO2 at 37°C.

### Antibodies, reagents and plasmids

The following primary antibodies were used: mouse anti-EEA1 monoclonal antibody (BD transduction, 610456), rabbit anti-VAPB polyclonal antibody (Atlas antibody HPA 013144), mouse anti-VAPA monoclonal antibody (Sigma 10355-4C12) and rabbit anti-VAPA polyclonal antibody for the immunoblot (Sigma SAB2702059), rabbit anti-TGN46 polyclonal antibody (abcam ab50595), mouse anti-paxilline monoclonal antibody (Merck, 05-417), mouse anti-cortactin monoclonal antibody (Merck, 05-180), mouse anti-tubulin monoclonal antibody (GeneTex GTX628802), mouse anti-PI(4,5)P2 monoclonal IgM antibody (Echelon Z-P045). The following secondary antibodies were used: goat anti-mouse IgM Alexa fluor 555 (Invitrogen, A21426) or Alexa fluor 488 (Invitrogen, A21042); goat anti-mouse IgG Alexa fluor 555 (Invitrogen, A21424) or Alexa fluor 488 (Invitrogen, A31620); goat anti-rabbit IgG Alexa fluor 555 (Invitrogen, A21429) or Alexa fluor 488 (Invitrogen, A11033).

The following reagents were used: fibronectin Hu Plasma Fibronectine EMD Milipore Corp; Methanol-free formaldehyde (Thermofisher, 28908); Fluoromont-G with DAPI (Invitrogen, 00-4959-52); Alexa fluor 647-conjugated Phalloidin (Invitrogen, A22287).

The following plasmids were used; pEGFP-MAPPER was a kind gift from Jen Liou (UT Southwestern Medical Center, USA) (Chang et al., 2013), pmCherry-Vinculin was a gift from Chinten James Lim (Addgene plasmid # 80024 ; http://n2t.net/addgene:80024 ; RRID:Addgene_80024) (Lee et al., 2013); pEGFP-PH-OSBP was a gift from Marci Scidmore (Addgene plasmid # 49571 ; http://n2t.net/addgene:49571 ; RRID:Addgene_49571) (Moorhead et al., 2010); pmRFP-PH-PLCδ1 was a kind gift from Francesca Giordano (Institute for Integrative Biology of the Cell, France) (Giordano et al., 2013); pmRFP-KDEL plasmid was a kind gift from Nihal Altan-Bonnet (National Heart Lung and Blood Institute, NIH, USA) (Altan-Bonnet et al., 2006); pmCherry-VAPA and pmCherry-VAPA KD/MD, obtained from pEGFP-VAPA and pEGFP-VAPA KD/MD, were kind gifts from Fabien Alpy (IGBMC, France) (Alpy et al., 2013); CRISPR-Cas9-resistant pmCherry-VAPA and pmCherry-VAPA KD/MD plasmids were obtained by mutagenesis of pmCherry-VAPA and pmCherry-VAPA KD/MD plasmids. Briefly, pmCherry-VAPA and pmCherry-VAPA KD/MD plasmids were amplified by PCR (30 cycles of 10s at 94°C, 5s at 55°C and 6min at 72°C) using the forward 5’TAAGACCGAATTCCGGTATCATCGATCCAGGGTCAACTGTGACTGTTTCAGT 3’ and reverse 5’ CGGAATTCGGTCTTACGCAATATCGTCGAGGTGCTGTAGTCTTCACTTTGA 3’ primers. PCR products were digested by DpnI enzyme for 2h at 37°C (Agilent) and loaded on a 1% Agarose Gel. PCR amplicons were extracted, purified using the NucleoSpin Gel and PCR clean up kit (Macherey-Nagel) and ligated using the NEBuilder HiFi DNA assembly kit (BioLabs).

### Transfection

For each transfection, 0.5 million Caco2 cells were nucleofected with 2 µg of DNA in 100uL of T solution using the B024 program on Amaxa Nuclefector II machine, as recommended by the manufacturer (Lonza). CaCo-2 transfected cells were then re-suspended in warm culture medium, replaced 24 hours later.

### Production of CRISPR Cas9 cell lines

Cells were transfected as described above with the double nickase CRISPR Cas9 plasmid (Santa Cruz, sc-424440-NIC) or a CRISPR Control plasmid (Santa Cruz, sc-437281). The following day, cells were incubated with puromycin at 2 µg/mL. After antibiotic selection, single GFP positive cells were sorted by flow cytometry. Clonal cells were maintained in culture with puromycin and the amount of VAPA protein was detected by western blot and immunofluorescence. VAPA KO cell lines expressing exogenous Cherry-VAPA and Cherry-VAPAKDMD were obtained by transfection of VAPA KO cells with CRISPR-Cas9-resistant pmCherry-VAPA and pmCherry-VAPA KD/MD plasmids. Stable cell lines were generated by selection with Geneticin (Gibco) and sorting of Cherry-positive populations by flow cytometry.

### Western Blotting

Confluent cells were lysed in 100 mM Tris pH 7.5,150 mM NaCl, 0.5% NP40, 0.5% triton-X100, 10% glycerol,1X protease inhibitor cocktail (Roche) and 1X phosphatase inhibitor (Phosphostop, Roche) for 20 minutes at 4°C. After 15 minutes centrifugation at 13000 g, solubilized proteins were recovered in the supernatant. Protein concentration was measured using Bradford assay (BioRad). For SDS PAGE, 50ug protein extracts were loaded in 4-12% Bis-Tris gel (Novex) or poly-acrylamide gels and proteins were transferred overnight at 4°C on a nitrocellulose membrane using a liquid transfer system (BioRad). Non-specific sites were blocked with 5% non-fat dry milk in PBS 0.1% Tween 20. Primary antibodies were diluted (1/1000 to 1/500) in PBS 0.1% Tween 20 and incubated overnight at 4ºC. After three washes in PBS 0.1% Tween 20, secondary HRP antibodies diluted in PBS 0.1% Tween 20 (1/10000) were incubated for 1 hour and washed 3 times with PBS 0.1% Tween 20. Immunocomplexes of interest were detected using Supersignal west femto maximum sensitivity substrate (ThermoFisher) and visualized with ChemiDoc chemoluminescence detection system (Biorad). Quantification of Western blots by densitometry was performed using the Gel analyzer plug in from Image J.

### Immunofluorescence

Cells were fixed with pre-warmed 4% formaldehyde in PBS for 15 min at RT and then washed 3 times with PBS. Permeabilization and blocking were performed in 0.05% saponin/0.2% BSA in PBS for 15 minutes at RT. The primary antibodies diluted (1/100) in Saponin/BSA buffer were then incubated overnight at 4°C. After 3 washes in saponin/BSA buffer, the samples were incubated with secondary antibodies (1/2000) and, when indicated, Alexa-coupled phalloidin (1/200) to stain F-Actin in the same buffer for 1 hour at RT. The preparations were washed twice in saponin/BSA buffer, once in PBS, and then mounted with the DAPI Fluoromount-G mounting media.

Immunostaining with anti-PI(4,5)P2 antibody was performed as described by Kim et al. (Kim et al., 2013). Briefly, cells were fixed at 4°C for 1h in 3.7% formaldehyde / 0.1% glutaraldehyde and incubated in PBS containing 0.1M glycine for 15 min. Permeabilization, blocking and staining were performed as described above, at 4°C.

### Cell migration assays

Collective migration assays were performed using 2-well silicon inserts (Ibidi). Glass coverslips were coated with fibronectin solution at 20 µg/mL in water, for 1h at room temperature. The cover slip surface was washed with sterile water and air-dried. Ibidi inserts were deposited on the fibronectin coated surface and 40000 to 50000 cells were loaded per well. For fluorescence live imaging, cells were directly plated after transfection in the well. After 3 to 4 hours of incubation, cell division was blocked using mitomycin at 10 µg/mL in CaCo-2 culture medium, for 1 hour. After overnight incubation in a fresh CaCo-2 culture medium, the insert was removed. Experiments were performed 24h to 48h after insert removal.

For individual cell migration, cell division was blocked using mitomycin at 10 µg/mL in CaCo-2 culture medium, for 1 hour. After overnight incubation in a fresh CaCo-2 culture medium, cells were detached and plated on a glass coverslip coated with 20 µg/mL fibronectin. After 6 hours, cells were imaged for 24h.

Collective and individual cell migration assays were performed using a Zeiss Wide-Field Microscope and imaged at 1 frame every 10 minutes. Cell trajectories were analysed using Manual Tracking from Fiji.

### Cell spreading assay

Cells were incubated for 1h on fibronectin-coated coverslips and fixed with formaldehyde 4% solution as described in the immuno labelling section. Coverslips were mounted on slides and samples were imaged using a Zeiss Apotome fluorescence microscope equipped with a 10x objective. The spreading area was determined based on Phalloidin-segmented ROI in Fiji/ImageJ.

### Nocodazole experiment

The nocodazole wash-out experiment was adapted from Ezratty et al (Ezratty et al., 2005). Briefly, cells were plated in the insert in starvation medium (DMEM GlutaMax medium containing 1% fetal bovine serum and 1% of penicillin/streptomycin). 24h later, the insert was removed, cells were left migrating for 24h in the starvation medium and treated with 10 µM nocodazole in starvation medium for 2h. Then, nocodazole was washed-out and cells were incubated with fresh starvation medium for 0, 15, 30 and 60 min and then fixed using 4% formaldehyde. Focal adhesions were labelled using a mouse anti-paxillin antibody as described above.

### Electron microscopy

Migrating cells were fixed in a 2% formaldehyde/1% glutaraldehyde in PBS solution for 1h at room temperature, then washed in PBS. Cells were embedded in a 10% BSA and 10% gelatin gel. Post fixation was performed in 1% OsO_4_+1% K_3_Fe(CN)_6_ in water at 4°C for 1h. After extensive washes, cells were incubated in 1% Thiocarbohydrazine in water for 20 min at room temperature and then washed and incubated with 2% OsO_4_ in water for 30 min at room temperature. After washes, cells were contrasted with 1% uranyl acetate overnight at 4°C. The following day, the uranyl acetate solution was removed and the samples washed using pure water. Cells were incubated with a pH 5.5 lead nitrate in aspartic acid solution for 30 min at room temperature, washed in water and dehydrated in successive ethanol baths. Cells were embedded in Agar Low Viscosity Resin (R1018 Kit from agar Scientific). 70 nm-thick thin sections were cut using a UC6 ultramicrotome from Leica and deposited on EM grids. Electron microscopy acquisitions were performed using a 120kV Tecnai 12 electron microscope (ThermoFisher) equiped with a OneView 4K camera (Gatan).

### Analysis of protrusive activity

The analysis of protrusive activity was performed on DIC images acquired every 3 seconds for 30 minutes. Kymographs representing the evolution of the leading edge in time along a line were generated and analysed with Fiji. Positive slopes were considered as protrusions and negative slopes as retractions. For each cell, the mean time spent in protrusion and the mean frequency of protrusions.was calculated from 2 to 4 kymographs.

### Measurement of focal adhesion assembly and disassembly

Cells were transfected as described above using the mCherry-Vinculine plasmid and migration assays were performed on fibronectin coated 1.5H Glass bottom Dishes (Ibidi). Cells were imaged using a TIRF microscope at 1 frame every minute for 1 to 2 hours. Focal adhesion assembly and disassembly rates were obtained using the focal adhesion analysis server (https://faas.bme.unc.edu/) (Berginski and Gomez, 2013). Focal adhesion assembly and disassembly rate tracks were generated. The tracks with a R-squared equal to or greater than 0.8 were used for the analysis.

### Quantification of FA in contact with ER-PM contact sites

The quantification of FA in contact with ER-PM contact sites was performed on TIRF microscopy acquisitions. 10 to 30 FA were randomly selected per cell. An FA was considered to be in contact with an ER-PM contact sites if there was at least one pixel recovery between GFP-MAPPER signal and mCherry-vinculin signal.

### Quantification of PI(4)P and PI(4,5)P2 levels

To characterize the intracellular pool of PI(4)P at the Golgi and on endosomes, colocalization between the GFP-PH-OSBP and TGN46 or EEA1 signals were obtained using Fiji Pearson coefficient plugins. The relative levels of PI(4)P in the Golgi or in endosomes were calculated as the ratio between GFP-PH-OSBP signal in the Golgi or endosomes masks and GFP-PH-OSBP signal in the cytosol, using segmentation tools in Fiji.

Concerning PI(4,5)P2, PM to cytosol signal ratio of RFP-PH-PLCδ1 was measured along lines of 5 to 15 microns at the leading edge of migrating cells, in protrusive area. The ratio was calculated as the maximal peak intensity along the line divided by the mean intensity value of RFP-PH-PLCδ1 in the cytosol. The plots on the graph represent the mean of 3 ratio at 3 different locations per cell.

For the quantification of endogenous PI(4,5)P2, cells were immnuno-stained with anti-PI(4,5)P2 antibodies. Confocal imaging was performed using the same acquisition parameters between control and VAPA KO cells. For one z-section, 3 lines were drawn per cell and the mean of maximal fluorescence intensity at the peak was plotted.

### Analysis of ER-PM contact sites on electron microscopy images

Electron microscopy images were analysed using Fiji/ImageJ. For the analysis, only ER-PM appositions of less than 30 nm distance were considered as ER-PM contact sites.

### Images acquisition and analysis

For live-microscopy experiments, the samples were placed in a chamber equilibrated at 37°C under 5% CO2 atmosphere.

Fluorescence live microscopy imaging was acquired using a Yokogawa CSU-X1 Spinning Disk confocal mounted on an inverted motorized Axio Observer Z1 (Zeiss) and equipped with a sCMOS PRIME 95 (photometrics) camera and a 63 X oil immersion objective or using an ELYRA TIRF microscope from Zeiss equipped with EMCCD, iXon 897 (512*512, 16 microns of pixel size) camera and 63 X oil immersion objective.

For fixed samples, images were acquired with a Zeiss Apotome fluorescence microscope equipped with a 63 X oil immersion objective or with a Zeiss LSM 780 confocal microscope equipped with a 63 X oil immersion objective at a resolution of 0.3 μm z-stacks. Image processing and analysis were done on Fiji software.

## AKNOWLEDGEMENTS

We thank Delphine Delacour, Fabien Alpy, Simon de Beco and the past and current members of Membrane Dynamics and Intracellular Trafficking team at Institut Jacques Monod for helpful discussions. We acknowledge the ImagoSeine core facility of the IJM, member of the France BioImaging infrastructure (ANR-10-INBS-04) and GIS-IBiSA, with support from La ligue contre le Cancer (R03/75-79), the Region Île-de-France (Sesame), Université Paris Cité (Labex Who Am I), Inserm (Plan Cancer), Région Ile de France (SESAME) and Fondation Bettencourt Schueller, France. This work was supported by grants from Gefluc groupement Les Entreprises Contre le Cancer, Cancéropôle Ile de France and INCa (2021-1-EMERG-51), La Ligue contre le cancer (RS21/75-85), the Agence Nationale de la Recherche (ANR-20-CE13-0021) and the Fondation pour la Recherche Médicale (DEQ. 20150934717), France. H.S was supported by fellowships from La Ligue contre le cancer and the Fondation pour la Recherche Médicale. M.L.H acknowledges René-Marc Mège, Benoît Ladoux and Delphine Delacour for kindly providing antibodies and plasmids during the starting period of this work.

## Author contributions

M.L.H. conceived the project. M.L.H and J-M.V supervised the project. H.S, R.LB, C.D, T.DA, A.V. and M.L.H performed experiments. M.L.H and H.S designed experiments. H.S, T.DA, A.V and M.L.H analysed data. M.L.H, H.S and J-M.V wrote the manuscript.

