## Supplemental Data for "The ER tether VAPA is required for proper cell motility and anchors ER-PM contact sites to focal adhesions"

### SUPPLEMENTAL FIGURE LEGENDS

**Supplemental Figure 1**

**A.** Left panel: representative immunoblots showing the levels of VAPB and Tubulin in Control and VAPA KO cells. Tubulin level was used as loading control. Right panel: quantification of relative VAPB density in Control and VAPA KO cells normalized to Tubulin levels (mean±SEM from three independent experiments). **B.** Confocal images of Control and VAPA KO leader cells immunostained for VAPB. Scale bar 20 µm. Data were analysed using non parametric Mann-Whitney t-test (ns: non significant).

**Supplemental Figure 2**

Confocal images of VAPA KO leader cells expressing wild-type mCherry-VAPA or mCherry-VAPA KD/MD. Scale bar 20 µm.

**Supplemental Figure 3**

**A.** Representative Transmission Electron Microscopy images of a Control and a VAPA KO cell showing ER-PM contact sites. Scale bar: 0.3 µm. The ER compartment and the PM are pictured in the right framebox. **B.C.** Analysis of the ER portion in contact with the PM (**B**) and the percentage of the PM in contact with the ER (**C**), quantified from images in A, in Control and VAPA KO leader cells (mean±SEM; Control: n= 25 cells: VAPA KO: n= 14 cells, from 3 independent experiments). All data were analysed using non parametric Mann-Whitney t-test (ns: non significant).

### SUPPLEMENTAL MOVIE LEGENDS

#### Supplemental Movie 1

Phase contrast movie showing collective migration behaviour of Control and VAPA KO cells 48h after space release. Scale bar: 100 µm.

#### Supplemental Movie 2

TIRF microscopy movie showing accumulation of GFP-MAPPER foci (green) at spots of close contact between the ER (Magenta) and the PM, at the front of a Control leader cell. Scale bar: 1 µm.

#### Supplemental Movie 3

TIRF microscopy movie showing the dynamics of GFP-MAPPER foci at the front of Control and VAPA KO leader cells expressing GFP-MAPPER. Scale bar: 1 µm.

#### Supplemental Movie 4

TIRF microscopy movie showing the dynamics of FA (Cherry-Vinculin, red) and GFP-MAPPER foci (green). The arrow points to the FA analysed in Fig.6D. Scale bar: 1 µm.

**Supplemental Figure 1**

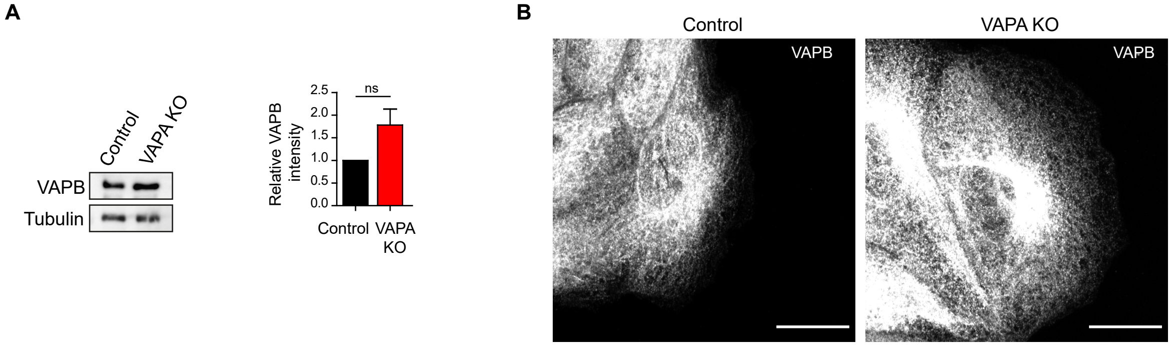

**Supplemental Figure 2**

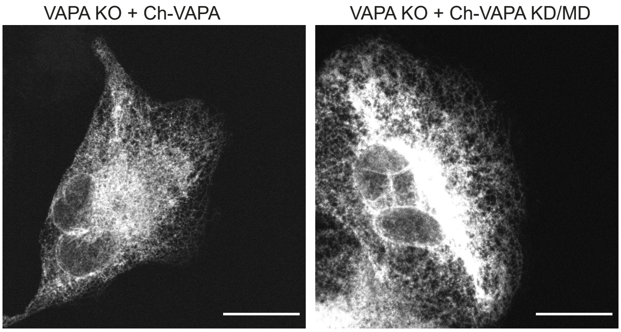

### Supplemental Figure 3

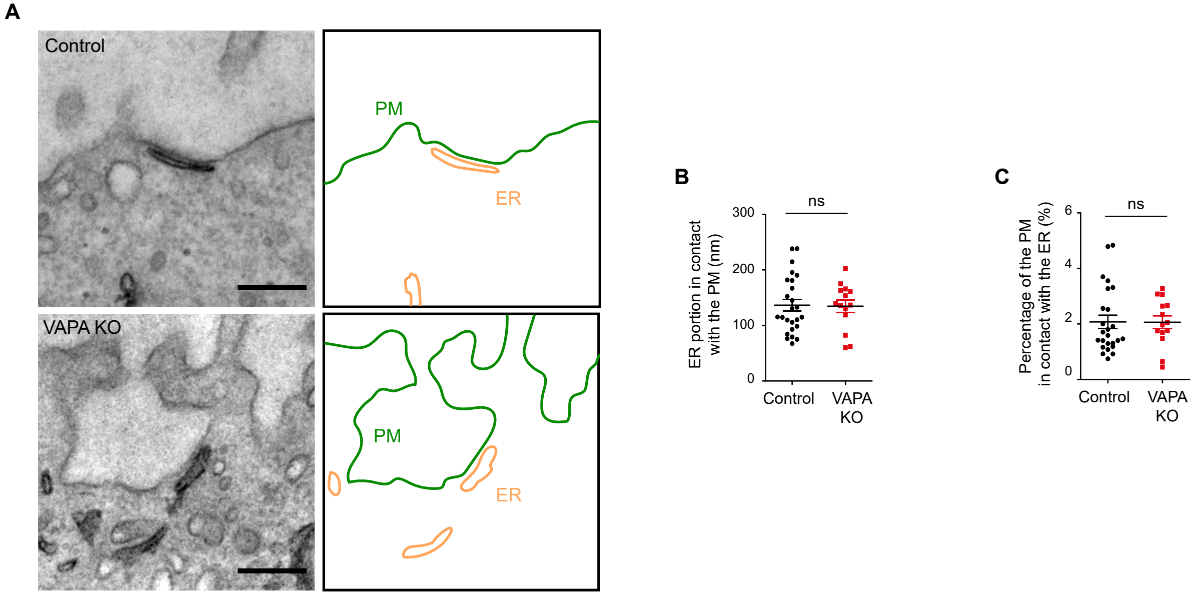
